# Comparing pest survey methods through detection probability: seasonal and abundance-dependent detection of spotted lanternfly (*Lycorma delicatula*, Hemiptera: Fulgoridae)

**DOI:** 10.64898/2026.10.29.762895

**Authors:** Joseph A. Keller, Matthew R. Helmus, Kelli Hoover

## Abstract

**BACKGROUND:** Invasive species management demands accurate information on the distribution of targeted pests, but surveys have imperfect detection. Understanding the factors influencing detection probability can guide improved management planning. For the invasive spotted lanternfly (*Lycorma delicatula*), several sampling methods have been used. We evaluated three widely applied methods: 5-minute timed visual search, BugBarrier sticky band traps, and modified circle traps. These methods were used to quantify abundance in 5 × 20 m plots spanning the core to margins of five outlying satellite populations in Pennsylvania U.S.A. Methods were assessed every two weeks over the growing season for two years.

**RESULTS:** Using a hierarchical Bayesian model, we found seasonal variation in all methods’ detection probabilities, with timed visual search increasing in sensitivity over the course of the season and both tree-based traps showing a U-shaped pattern, with lowest detection probability around midseason. For timed visual search in plots used by ten *L. delicatula* individuals, the estimated probability of detecting at least one individual ranged from 10% to 53%. For tree-based traps, detection was higher on larger trees and on *Ailanthus altissima*, a preferred hostplant. For traps installed on *A. altissima* with diameter at breast height of 20 cm in a plot with the same *L. delicatula* density (N=10), estimated mean detection probability ranged from 33% to 99% for BugBarrier and from 28% to 84% for circle traps.

**CONCLUSION:** Informed with estimated detection probabilities, managers can compare alternative survey methods and design strategic management plans that properly incorporate realistic survey sensitivity.

## 1. Introduction

Detection, the process by which an organism is documented at a location where it is present, plays an important role in efforts to limit the negative impacts of invasive species.^1^ Initial documentation of new invasive species depends on detection,^2^ and biosecurity measures aim to proactively detect potential pests in transported goods.^3^ Further, surveillance around points of entry aims to shorten the timespan between pests’ introduction and discovery to catch emerging pests before they spread.^4^ Once invasive pests are established, surveys to locate their distribution are critical to guide management.^5^ Delimiting surveys have been widely applied for pests such as the spongy moth (*Lymantria dispar*)^6^ and Asian longhorned beetle (*Anoplophora glabripennis*)^7^. Effective detection processes remain valuable for widespread, long-established invasive species. Efforts to track population dynamics,^8^ evaluate infilling,^9^ and measure management impacts^10^ hinge on detection effectiveness. Finally, when attempts to eradicate invasive species are made, detection of potential remnant individuals is necessary to evaluate outcomes.^11^

There are a variety of methods available to managers surveying for invasive pests, each with varying detection probabilities. Visual surveys ^12^, traps with ^13^ and without attractants,^14^ environmental DNA (eDNA) collection,^15^ remote sensing,^16^ citizen science programs,^17^ acoustic surveys,^18^ and detector dog surveys^19,20^ all are used to detect invasive species. For each of these methods, detection probability may be influenced by their application (including observer effects),^21^ location,^22^ and timing.^23,24^ In practice, the methods selected depend on the biology of the pest, the availability of workers and materials, and the cost of deployment.^1^

The spotted lanternfly (*Lycorma delicatula*, Figure 1) is an invasive species with economic impacts on grapevines,^25–27^ and management to limit its spread hinges in part on managers’ ability to detect this pest.^28^ *Lycorma delicatula* feeds on the phloem of grapevines and can reduce yield and grape quality, as well as cause vine mortality.^27,29^ Its spread in the United States has been marked by repeated establishment of outlying satellite populations,^30–32^ which begin with low densities that can be difficult to detect. A variety of survey methods have been developed targeting *L. delicatula*. Dogs trained to detect *L. delicatula* by smell have been used to investigate potentially contaminated goods and facilities.^19^ Collection and analysis of eDNA has proven highly sensitive.^33–35^ A public awareness campaign encouraging people to report sightings of this distinctive insect has yielded hundreds of thousands of observations.^29,36^ Traps including sticky bands and circle traps exploit spotted lanternflies’ propensity for climbing trees and have been used for early detection and to quantify local densities.^37,38^ The toolset available to managers is diverse, but comparisons between methods are incomplete.

**Figure 1.**
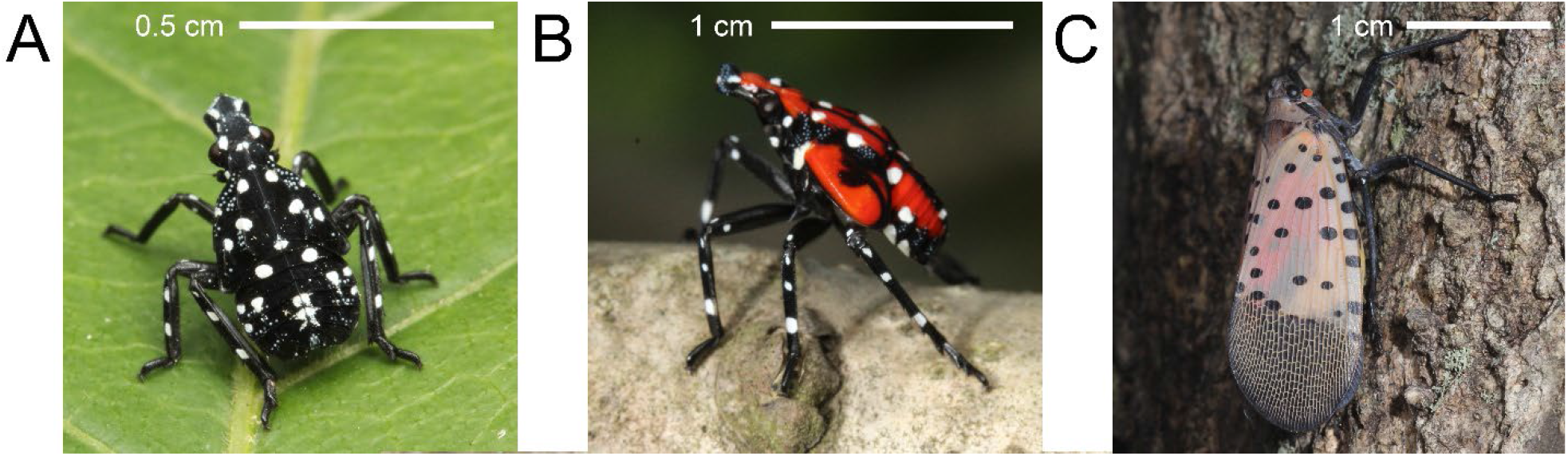
Spotted lanternfly individuals vary in conspicuousness over the course of their development. (A) Early instar nymphs (instars 1-3, second instar shown) have black bodies marked with contrasting white spots. (B) Fourth instar nymphs have a bright red patterning with black portions and contrasting white spots. (C) Adult forewings are pale greyish with black markings, while hindwings, which are displayed in response to harassment, are red, white and black. Scale bars are approximate.

We evaluated affordable and widely applied survey methods for detecting spotted lanternflies, estimating their probability of detection throughout the season. We tested the sensitivity of timed visual search, tree-based circle traps, and BugBarrier sticky bands across a range of *L. delicatula* densities and throughout the season. Based on two years of sampling data from outlying *L. delicatula* populations, we address the following questions:

1. How do the detection probabilities of these three sampling methods compare to one another?
2. How does *L. delicatula* abundance affect detection probability?
3. How do detection probabilities vary over the season?
4. For tree-based traps, how do tree size and tree species affect detection probability?

As *L. delicatula* continues to spread, better understanding of how reliably these widely used survey methods detect populations can improve surveillance and management outcomes.

## 2. Materials and Methods

### 2.1 Study Sites

We tested survey methods for detecting *L. delicatula* at sites around nascent outlying populations (Figure 2), allowing us to visit locations with varying *L. delicatula* densities within a compact area. In Pennsylvania in spring 2020, we established 28 20 x 5m plots in Altoona, 14 in Shippensburg, and 5 in Huntingdon. In spring 2021, four new sites were added in Altoona (and one was lost due to revoked access), two were added in Shippensburg, six were added in Huntingdon, and seven were added in each of Lewistown and Johnstown, PA. Plots were located along forest edges and in strips of marginal, minimally managed vegetation. They were generally oriented with the long edge along the forest edge or vegetation strip. All plots contained host plants for *L. delicatula*,^39^ and most, though not all, plots contained *Ailanthus altissima* (58 of 73 plots), one of *L. delicatula*’s preferred hosts.^40^

**Figure 2.**
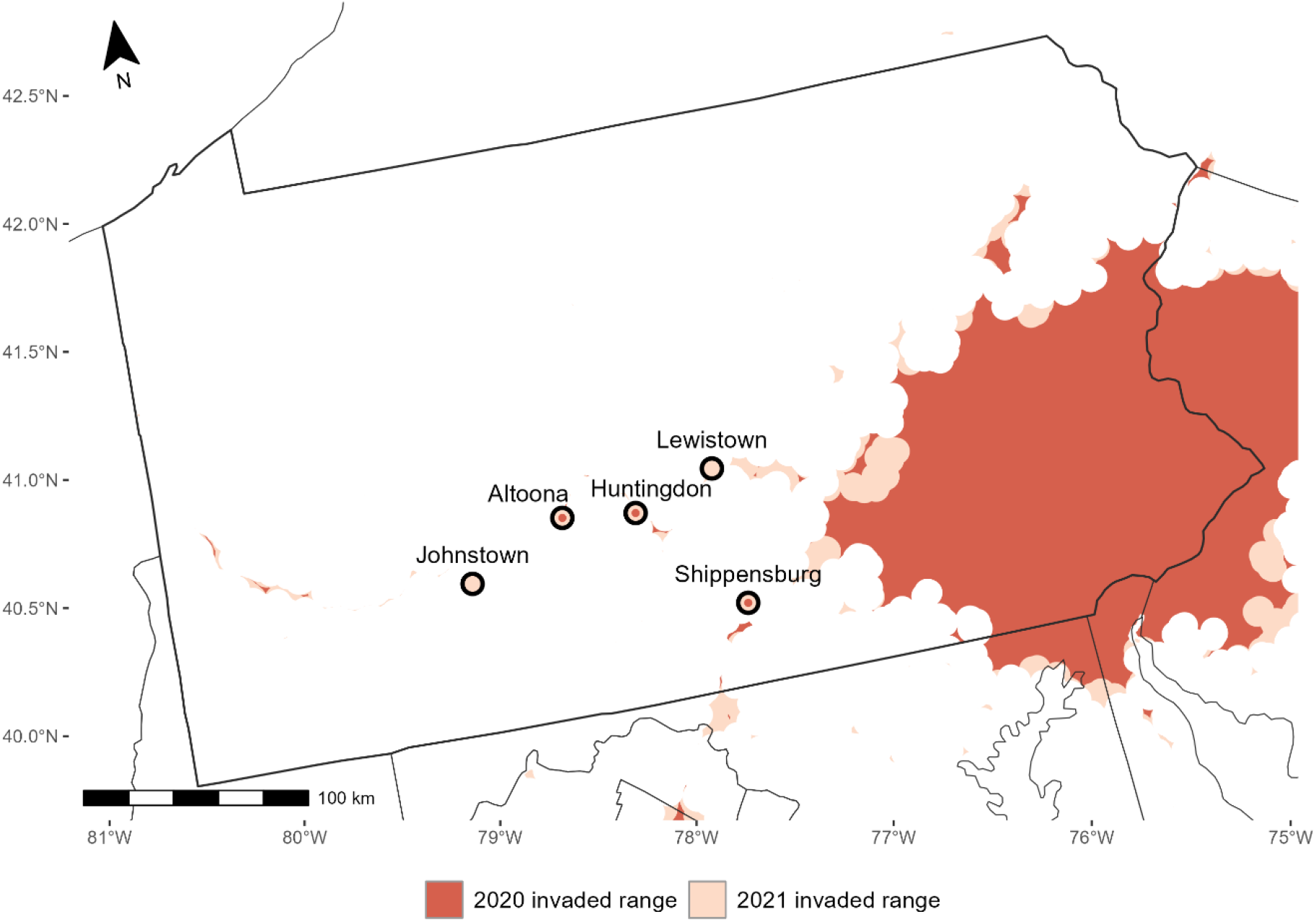
Sampling methods were tested at outlying *Lycorma delicatula* populations in Pennsylvania in 2020 and 2021. Cities where sampling occurred are marked with points, and the extent of the invasion is shown for 2020 and 2021 (bounded by alpha convex hulls around presences reported by state and federal surveyors^32,36^).

### 2.2 Sampling methods

We tested three sampling methods: timed visual search, inward-facing sticky BugBarrier traps, and modified circle traps (Figure 1). For each of these methods, the number of *L. delicatula* captured or seen was counted. Plots were visited every other week from May through November. In the first year of observation, BugBarrier was installed in early July, and circle traps were installed later, in mid to late August. In 2021, both traps were installed at the start of the season. Within each plot, we swapped the trap type between trees on the first visits following 29 June 2021 and 27 September 2021 to avoid confounding trap type with tree attractiveness, which is known to vary.^41^

To carry out timed visual search, we approached the edge of the plot and started a five-minute timer. For the duration, two observers moved through the plot counting all *L. delicatula* nymphs and adults that they could see (Figure 1). Observers were instructed to cover the entire plot and were permitted to revisit areas as they pleased during observation period. If necessary, observers moved vegetation, but care was taken not to disturb any individuals discovered. No binoculars or other aids were used. On each visit to the plot, two observers conducted independent timed counts simultaneously, except rare instances when labor was unavailable and only one count was made.

We installed one modified circle trap per plot (Figure 3A). These traps were pecan weevil trunk traps (GL/GL-4010-06, Great Lakes IPM, Vestaburg, MI) modified to capture spotted lanternflies. The hole in the upward-facing funnel at the top of the trap was widened to have a diameter of approximately 1.5 cm as in Francese et al. (2020). To contain captured individuals, we modified the plastic cup container by removing its closed end, hot gluing a u-shaped portion of plastic cup to extend one edge of the resulting cylinder, putting a ring of hot glue around the outside of the container, and zip-tying a plastic bag in place. The addition of the u-shaped plastic extension ensured the bag did not fall flush with the end of the original container and block individuals from entering the bag. The inside of the plastic cylinder was lightly sanded to ensure spotted lanternflies could climb over this vertical surface. Traps were installed on trees using staples, with the bottom edge of the mesh pulled snug against the trunk. Traps were installed at approximately chest level on preferred hosts of spotted lanternflies in the following order of priority based on availability within the plot: tree of heaven (*Ailanthus altissima*, chosen first if available), red maple (*Acer rubrum*), black cherry (*Prunus serotina*), black walnut (*Juglans nigra*) and sumac (*Rhus sp*., chosen last if no other hosts were available). Neither chemical attractants nor insecticides were included with the trap. On each check of these traps, the bag was inspected for spotted lanternflies. If any were found, the bag was removed and replaced. If no individuals were found, the funnel opening was checked for spiderwebs, cleared if present, and the bag was left in place.

**Figure 3.**
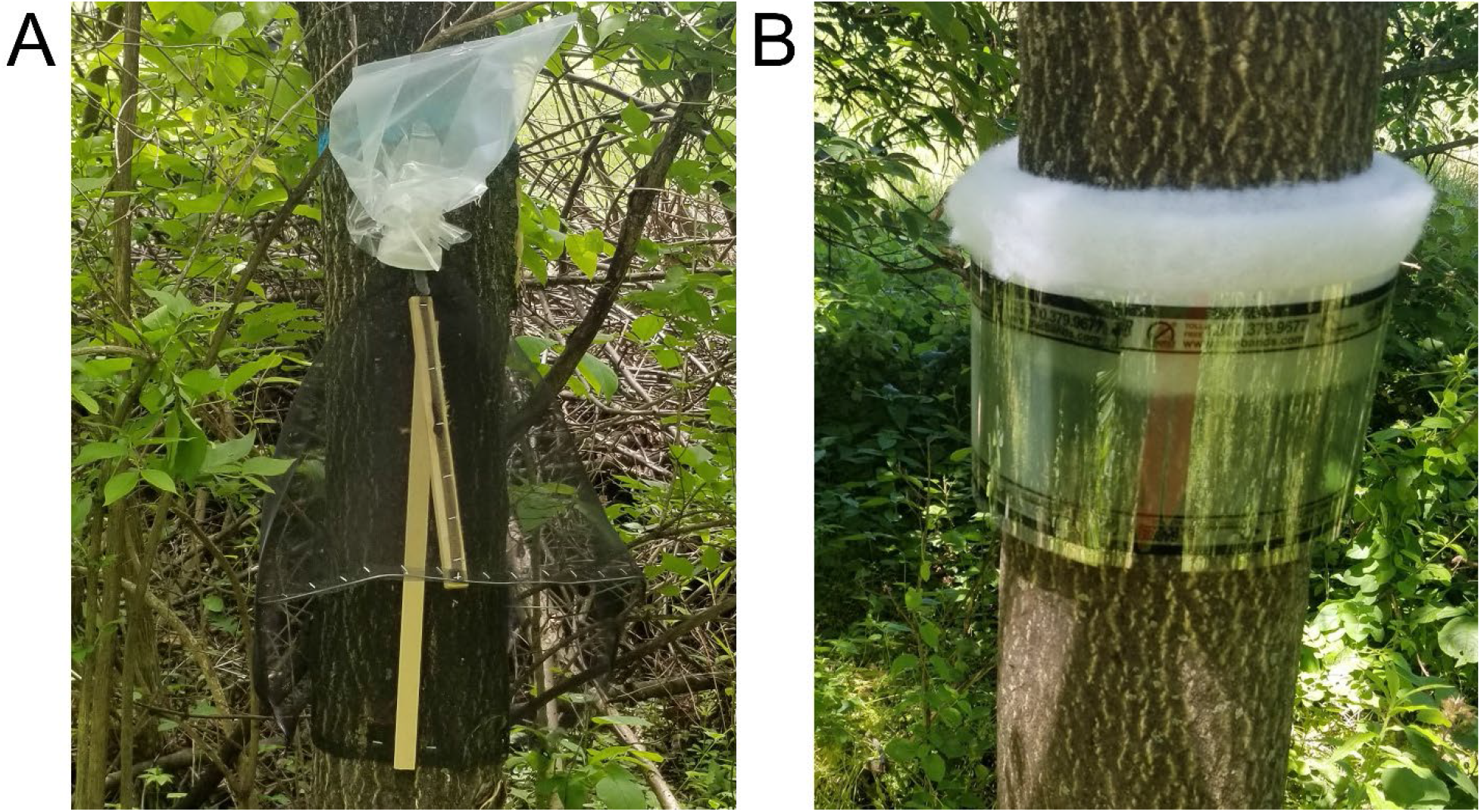
Tree-based traps capture spotted lanternflies (*Lycorma delicatula*) by exploiting their climbing behavior. Circle traps (A), which are modified from pecan weevil traps, use netting around the trunk to funnel climbing individuals into a retaining bag. BugBarrier sticky bands (B) use a fiber material wrapped around tree trunks to achieve a gap between the trunk and an inward-facing adhesive film.

We deployed inward-facing BugBarrier sticky band traps on one tree per plot (Figure 3B). Following the same host plant preference order, we wrapped two layers of fiber material around the trunk of the selected tree. We then wrapped the BugBarrier sticky film around the fiber with the adhesive side facing inward, leaving between half and two thirds of the 12.7 cm-width surface exposed. On each check of the BugBarrier sticky bands, the sticky plastic was removed and inspected for spotted lanternflies. Fiber material was replaced as needed to maintain sufficient spacing from the trunk for lanternflies to move past the bottom rim.

### 2.3 Growing Degree Days

To characterize the seasonal timing of observations, we calculated accumulated growing degree days for each outlying population. We used weather station data from the Daily Global Historical Climatology Network.^42^ Data from the following weather stations were used: USC00364992 (Lewistown), US1PAHN0003 (Huntingdon), USC00360140 (Altoona), USC00368073 (Shippensburg), and US1PACM0004 (Johnstown). We calculated degree days using the sine method with lower threshold 10.4°C and upper threshold 99°C using the *dd_sng_sine* function from the *degday* package in R.^43^ The lower threshold temperature was based on prior developmental and modeling work,^44,45^ and the upper threshold was set above observed temperatures.

### 2.4 Statistical analysis

We constructed a hierarchical Bayesian model to assess the detection probabilities of timed visual search, circle traps, and BugBarrier sticky band traps. We evaluated candidate models containing different combinations of the available predictors (Table S1, S2, Figure S1), but here we describe only the final selected model. Predictor variables were centered and scaled. In all models, each plot-year combination was treated as a distinct site. This structure was appropriate because spotted lanternfly abundance can change substantially at the same plot between years as the invading populations expand or shift locally, while repeated surveys within a year sample the same underlying local population. Under this structure, we had 119 plot-year sites, with sampling divided into 12 two-week intervals for each site.

Because the survey methods operated over different observation periods, we defined abundance as a superpopulation.^46,47^ In this model, the superpopulation was the total number of individuals that used the site at some point during a given approximately two-week interval, rather than the number present in the site at any single moment. The traps set for two-weeks could therefore capture individuals from throughout this superpopulation as they moved through the plot during that interval, whereas a five-minute visual search sampled only those members of the superpopulation that were present during the brief search period. For invasive species surveys, particularly at low-density sites, this superpopulation perspective aligns with management program priorities because identifying locations invaded by the targeted pest is often more important than determining whether individuals are present within the surveyed area at a single point in time. Consequently, the estimated detection probability for visual search captures two processes: the probability that an individual in the two-week superpopulation was physically available within the site during the search and the probability that an available individual was successfully detected by the observer.

Within this superpopulation framework, we modeled the underlying ecological process in two stages: first assessing whether a site was occupied and, conditional on occupancy, tracking how abundance changed through time. Specifically, site occupancy was modeled using a dynamic process with parameters for the rate of colonization in unoccupied sites (*γ*) and the rate of persistence in occupied sites (*φ*). For each site *i*, initial occupancy (*Z*_*i*,1_) was modeled as a Bernoulli process (a yes/no outcome with a fixed probability):

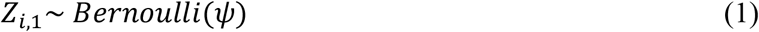

where *ψ*is the initial occupancy probability. From there, occupancy in each site progressed though colonization of previously unoccupied sites and persistence of occupied sites.

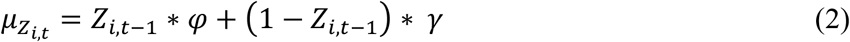

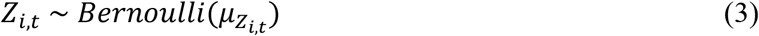

In this model, colonization probability was held constant across sampling intervals. We also evaluated a model with time-varying colonization, but the additional interval-specific colonization parameters did not converge despite extended model runs, so we retained this simpler constant-colonization structure.

Within occupied sites, we modeled potential abundance following a temporally autocorrelated random walk. At each time step, superpopulation abundance *N* followed a Poisson distribution (a random count with a given average) constrained by site occupancy.

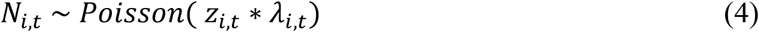

The parameter *λ*_*i,t*_ *wa*s initialized with a site level intercept *α*_*i*_:

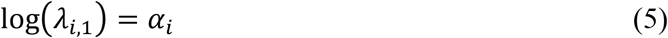

and progressed with Gaussian process error *ω*_*i,t*_.

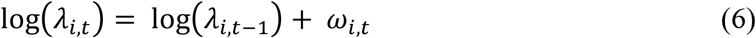

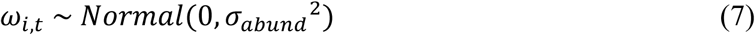

The process variance *σ*_*abund*_^2^ captured the magnitude of fluctuations in abundance on the log scale.^48^

Counts that were made during timed visual search were modeled using a beta-binomial distribution with abundance-dependent mean detection probability:

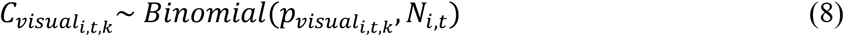

where *k* denotes which observer made the count and 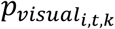 *is* the probability of detection by observer *k* in site *i* at time *t*. This probability was modeled using the beta-binomial distribution to account for overdispersion after simpler binomial models showed poor goodness of fit ^49^:

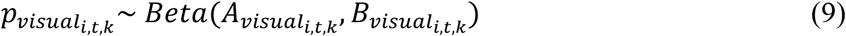

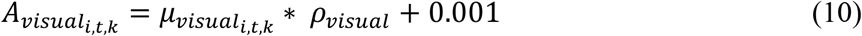

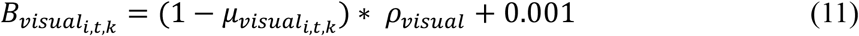

where *ρ*_*visual*_ controls the level of overdispersion. A small constant (0.001) was added to both shape parameters to ensure *A*_*visual*_ and *B*_*visual*_ remain strictly positive to facilitate Markov Chain Monte Carlo sampling. The mean detection probability 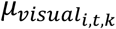 was modeled as follows:

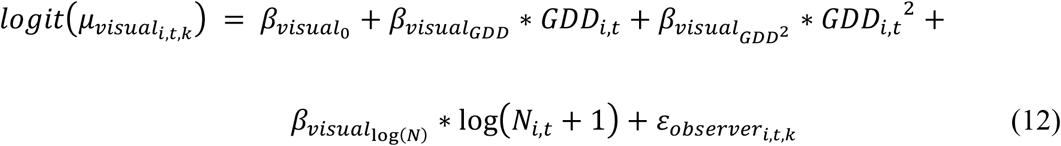

Seasonal patterns in detection probability were captured with 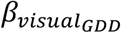 *wh*ich described the effect of accumulated growing degree days (*GDD*_*i,t*_) and in the squared term 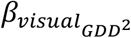 .

Potential variation in detection related to abundance in the plot was described using 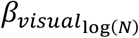. A positive fitted value for this parameter would indicate better detection when greater numbers of individuals were present in the plot, as might be expected if individuals were aggregated in clusters as has previously been observed for this species.^50^ Systematic differences between observers were characterized with the effect 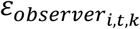 with 6 individual observers included in the dataset and these effects constrained to sum to 0.

For each of the two tree-based traps, we used identical model structure with method-specific parameters. Trap catch counts were modeled as beta-binomial draws from the superpopulation:

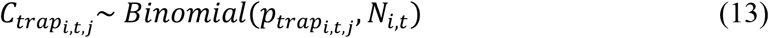

where *j* denotes which of the two traps was used. Similarly to the assessment of visual counts, we used the beta-binomial distribution to account for overdispersion in trap catch.

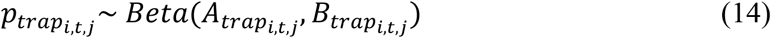

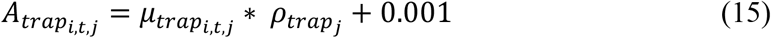

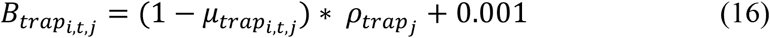

Here, 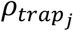 *co*ntrolled the amount of overdispersion for trap type *j*. The mean detection probability for traps was modeled as follows:

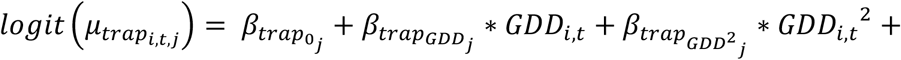

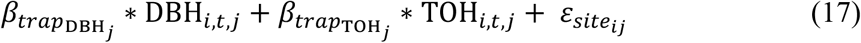

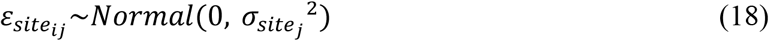

The site-level random effect 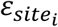 *wa*s included to account for site-level variation in trap effectiveness, as trees are known to have varying levels of use by *L. delicatula*.^41^ Variation between sites was quantified in 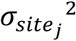, the variance associated with this random effect. The diameter at breast height (DBH) of the tree on which traps were installed was included to account for the influence of tree size. An indicator variable ‘TOH’ documented whether the trap was installed on a tree-of-heaven (*A. altissima*) tree (1) or on another species (0). As in the visual detection model, seasonal variation in detection probability was modeled with the linear and squared GDD effects.

To fit the model, we employed JAGS ^51^ implemented using the *jagsUI* package in R.^52^ We used three parallel MCMC chains with a 10,000 adaptation phase (discarded) followed by 350,000 sampling iterations, of which the first 50,000 were discarded as burn-in. Sampling was thinned by taking one value every 35 iterations. Model convergence was assessed based on the Gelman-Rubin metric 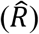, and we ensured the final fitted model had all 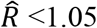 *f*or all monitored parameters.^53^ We additionally assessed the goodness of fit for each survey method’s detection model based on the Freeman-Tukey discrepancy measure evaluated at the level of individual detection records.^54^ We considered posterior predictive *p*-values between 0.1 and 0.9 to indicate adequate fit. We used uninformative priors for all parameter classes except for three that we assigned moderately informative priors. For the initial abundance intercepts *α*_*i*_ we used a moderately informative prior to constrain abundances to a biologically plausible range while maintaining flexibility (Table S3).^55^ For the visual detection overdispersion parameter *ρ*_*visual*_, we used a moderately informative prior to prevent extreme variability in detection based on the beta-binomial formulation (Table S3). And for the observer effects on visual detection, we used a moderately informative prior (SD = 2 on the logit scale) to stabilize estimation of observer effects that were weakly informed arising from unbalanced observer sampling effort. For a full list of model parameters, and their priors, see Table S3. Additionally, to identify potential trade-offs among parameters we evaluated correlation between model parameters’ posterior samples. Finally, we assessed parameter identifiability by using data-cloning procedures with 1, 3, and 6 clones of the data.^56^

## 3. Results

Sites had a range of *L. delicatula* abundances, with observed counts ranging from 0 to 1031. Duplicate timed visual counts were generally close to one another, though variation in counts indicated imperfect detection, as expected (Figure 4). Over the course of the study, we made 1070 observations with the complete set of all four surveys (both timed visual searches, BugBarrier trap, and circle trap). Cases with missing values were caused by occasional staffing absences and damage to traps including lost bags from circle traps and animals (e.g., squirrels) removing fiber material from BugBarrier traps. The most common detection pattern was no detection by any of the four surveys (n=421, Figure 4), likely owing to plots’ locations around small, newly discovered *L. delicatula* populations with low abundance and limited spatial extent. Second most common (n = 277) were cases where all four surveys were positive.

**Figure 4.**
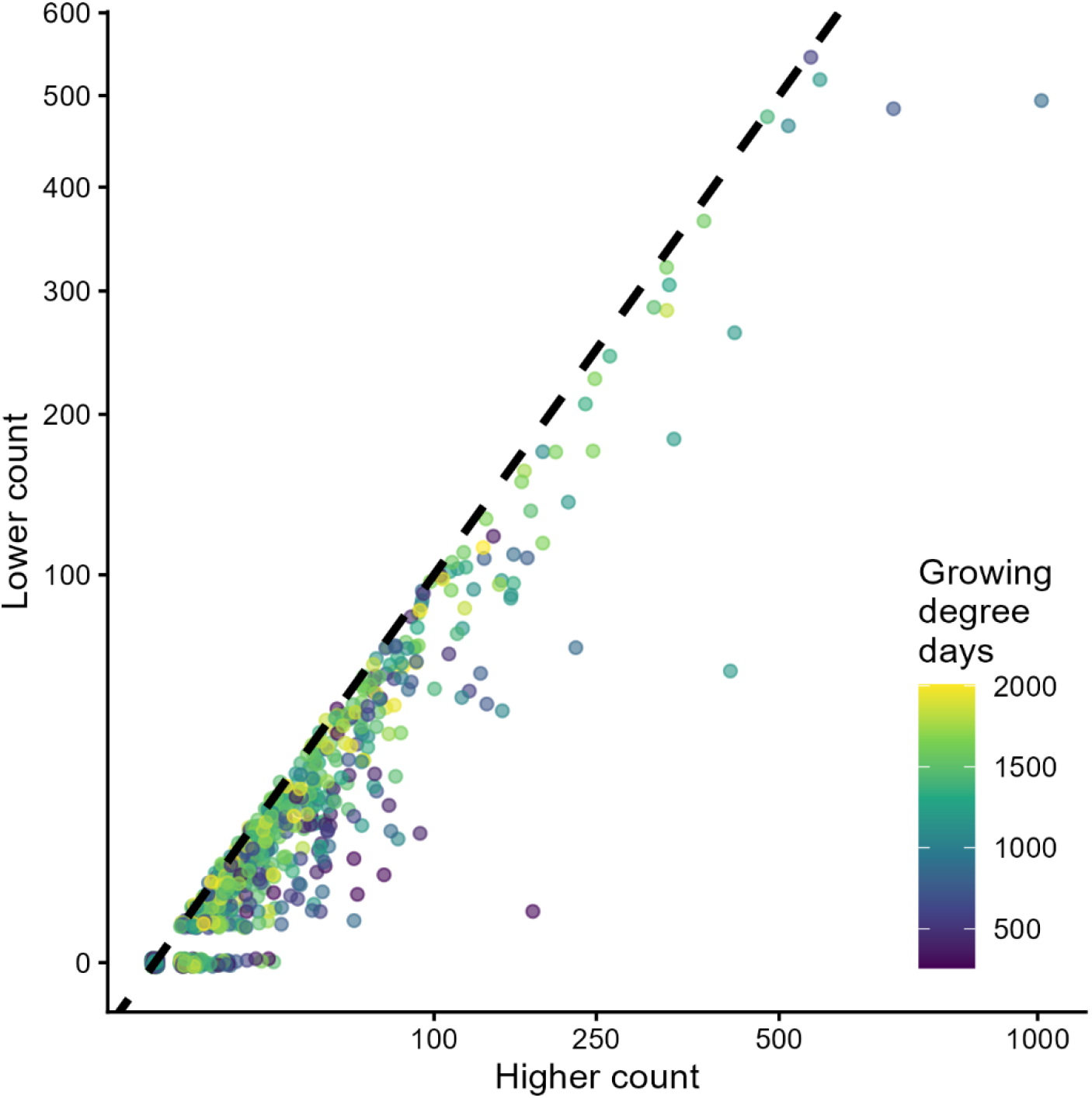
Duplicate 5-minute visual surveys generally produced similar counts (points falling near the 1:1 dashed line), but disagreement between paired observers provided direct evidence of imperfect detection and the need to account for observation error when estimating *L. delicatula* abundance. Points are colored based on the accumulated growing degree days that had accrued by the time counts were made. Both axes are square-root transformed to better show the bulk of the data with lower numbers of individuals. Points were also randomly jittered slightly (within 0.1 of true values) to reduce overlap.

The fitted model indicated a strong seasonal trend in the detection probability of timed visual search (Figure 5). Early in the season, when small, black-and-white nymphs were present, the estimated probability of detecting a single individual from the two-week superpopulation was around 1% (at GDD=400, 95% CI [0.04%, 1.4%]. The average probability detecting at least one individual increased with superpopulation size. For example, a superpopulation of 10 individuals using the 100 m^2^ site over the 2-week period (equivalent to 0.1 individuals m^-2^) had an estimated 15% probability of at least one detection at GDD=400 (95% CI [9%, 23%]). Detection improved later in the season as larger adults became prevalent. At GDD=1900, the estimated probability of detecting a single individual was 4% (95% CI [2.2%, 6.7%]), while the probability of detecting at least one individual from a superpopulation of 10 was 54% (95% CI [38%, 71%]). For the visual detection portion of the model, goodness of fit as measured using the Freeman-Tukey discrepancy metric was acceptable (Bayesian p-value = 0.55, figure S2).

**Figure 5.**
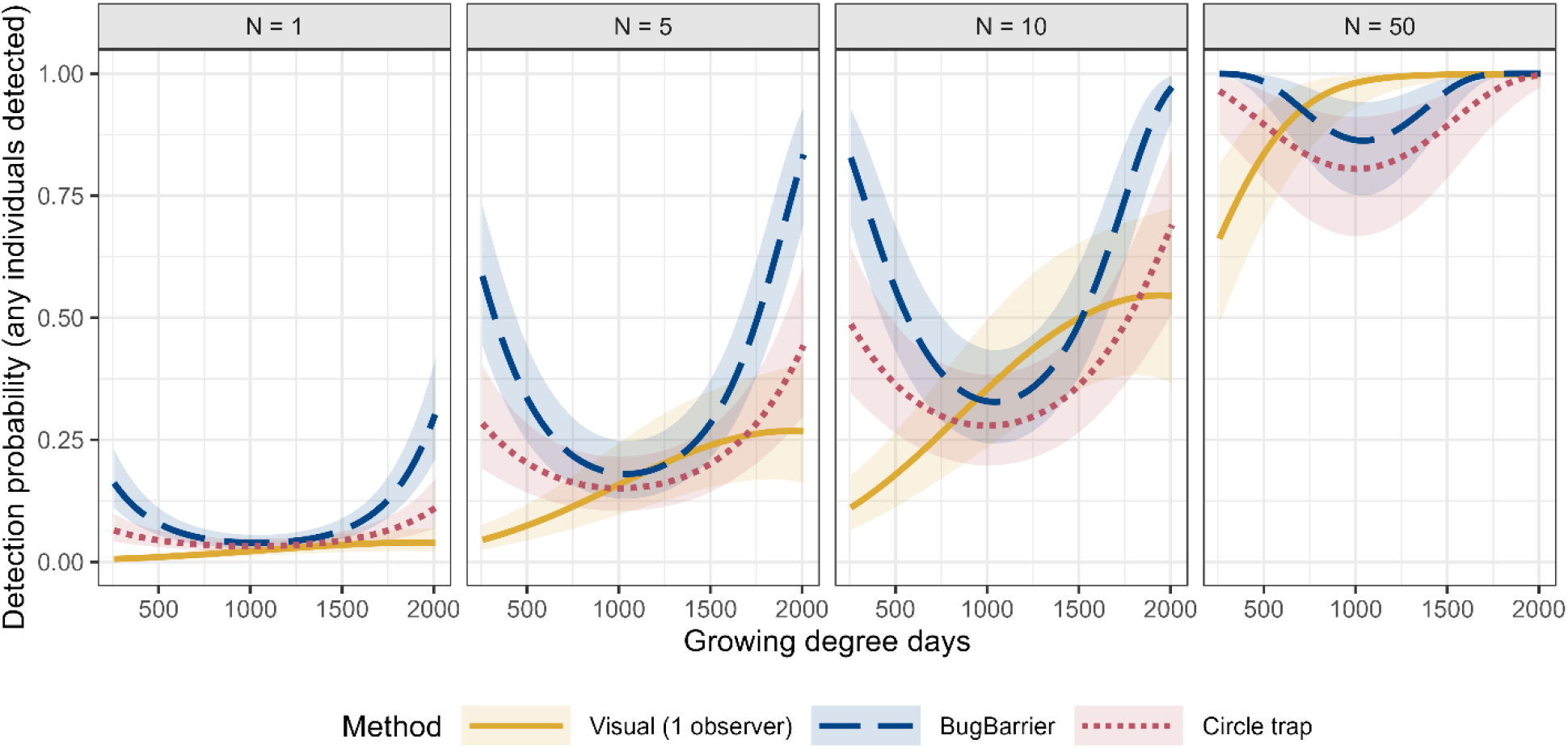
Model-estimated detection probabilities showed that survey effectiveness depended on both *L. delicatula* abundance and seasonal timing. The probability of detecting one or more spotted lanternfly individuals increased with increasing superpopulation size (compare across facets) and varied by method (compare different colored lines) and across the growing season (compare along the x-axis). Detection by timed visual search (yellow) increased over the course of the season, while detection by both tree-based traps started high, fell mid-season, and improved again late season (note the u-shaped curves for BugBarrier, blue, and circle traps, red). At low *L. delicatula* densities, mean detection probabilities were estimated to be low (first panel, superpopulation size N=1), falling under 20% for all methods for much of the season. At high *L. delicatula* densities (last panel, superpopulation size N=50), timed visual search performed very well, with near certain detection after mid-season. For tree-based traps, fitted lines show estimated mean values for installation on *Ailanthus altissima* with diameter at breast height 20 cm.

For BugBarrier sticky band traps, the probability of detecting at least one individual similarly increased with superpopulation size and also varied with tree size, tree species and seasonal timing (Figure 5). Per-individual detection probability increased with the diameter of the tree on which the trap was installed (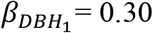, 95% CI [0.20, 0.42]) and was higher for traps installed on *A. altissima* (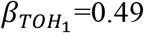, 95% CI [0.15, 0.84]) than on other tree species. For example, at GDD = 1500 (around the onset of oviposition) a BugBarrier trap installed on a 10 cm DBH non-*A. altissima* tree had an estimated 22% chance of detecting at least one individual from a superpopulation of 10 individuals (95% CI [15%, 31%]). Under the same conditions, a trap installed on a 30 cm DBH *A. altissima* tree had a 67% probability of detecting at least one individual (95% CI [52%, 81%]). Detection also varied strongly over the course of the season, declining from relatively high levels early in the season to a minimum at GDD = 1041 when fourth instars were abundant, before increasing again later in the season when adults were mating and laying eggs. Both the linear and the squared terms for accumulated growing degree days were substantially different from zero 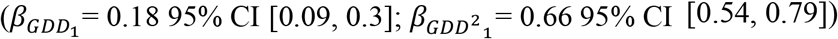. The BugBarrier portion of the model showed satisfactory convergence (Rhat < 1.05) and acceptable goodness of fit (Bayesian p-value = 0.38, Figure S2).

Circle trap detection probability followed similar patterns to those observed for BugBarrier (Figure 5). Per-individual detection probability was higher on larger trees (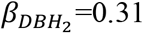, 95% CI [0.18, 0.45]). Detection probability did not differ substantially between traps installed on *Ailanthus altissima* and other species; the 95% credible interval for this effect included zero (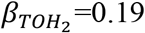, 95% CI [-0.22, 0.60]). Detection varied strongly over the growing season. Like for BugBarrier traps, circle trap per-individual detection probability followed a U-shaped pattern, with a minimum at GDD =1001 and higher detection earlier and later in the season (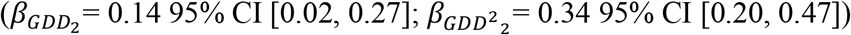. The circle trap portion of the model showed satisfactory convergence (Rhat < 1.05) and acceptable posterior predictive fit (*p*-value = 0.89, Figure S2)

The survey method with the highest probability of detecting at least one individual depended on seasonal timing, superpopulation size, and the size and species of the tree available for trap installation (Figure 6, Figures S3, S4). Early and late in the season, BugBarrier generally had the highest estimated detection probability. During the middle of the season, timed visual search outperformed both tree-based traps. However, when large *A. altissima* were available for trap installation, BugBarrier remained the best-performing method across much of the season, particularly at low superpopulation sizes. Under these conditions, timed visual search exceeded BugBarrier primarily at higher *L. delicatula* superpopulation sizes, where the probability of detecting at least one individual was relatively high regardless of the method used. Circle traps outperformed the other methods at low *L. delicatula* superpopulation size at the midseason, but only when traps were installed on large, non*-Ailanthus* trees. Differences among methods were relatively small under these conditions (Figure S4).

**Figure 6.**
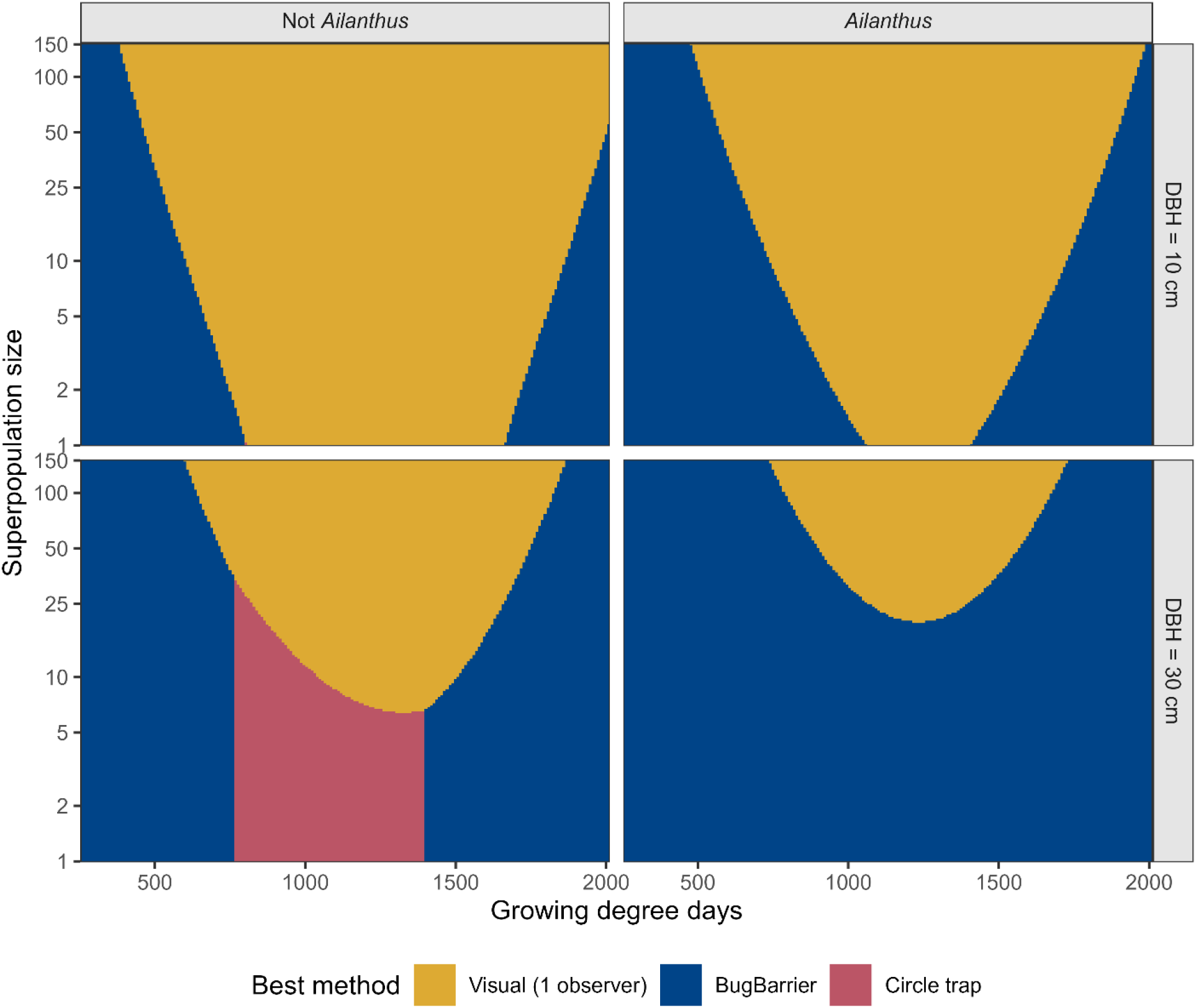
No single survey method consistently provided the highest detection probability. The choice of the best performing survey method was influenced by the species of the tree on which traps were installed (columns), the size of the tree on which traps were installed (rows), the seasonal timing of observation (x axis in each phase plot) and the superpopulation size (y axis in each phase plot). Areas within each panel are colored to identify which sampling approach had the highest probability of detecting one or more individuals under the indicated conditions. When the best tree available for trap installation is a small, non-Ailanthus tree (top left panel), timed visual search outperforms both tree-based traps for much of the season and across a substantial range of *L. delicatula* densities (note the large yellow area). When there is a large *A. altissima* tree available (bottom right panel), BugBarrier trapping outperformed timed visual search and circle traps over the entire season at low *L. delicatula* densities (note the blue color spanning the x-axis for low superpopulation sizes).

Posterior samples showed substantial correlation among some detection and abundance parameters. Within each beta-binomial detection model, the detection intercept was negatively correlated with the concentration parameter (-0.71 to -0.56). This correlation is expected because the mean detection probability and the beta-binomial concentration parameter (ρ) jointly determine the distribution of observed counts, allowing some combinations of these parameters to produce similar predictions. We additionally observed positive correlations among corresponding parameters across the three detection models: detection intercepts were correlated across methods (*r* = 0.37 to 0.53), as were concentration parameters (*r* = 0.60 to 0.61). These cross-method correlations likely arose because all three observation models were linked through the shared latent abundance *N*, such that changes in estimated abundance produced similar changes to detection parameters across methods. These posterior correlations indicate parameter trade-offs during estimation but do not, by themselves, imply non-identifiability. Data cloning analysis with *K =* 1,3,6 copies of the dataset showed that posterior variances for all detection parameters decreased approximately in proportion to 1/K, demonstrating that the likelihood surface possessed a well-defined global maximum and that these parameters were identifiable under the observed data structure (Table S4).

## 4. Discussion and Conclusions

By co-deploying three commonly used survey methods for the invasive pest *L. delicatula*, we found that survey effectiveness varied substantially across the growing season and depended on both local abundance and the characteristics of trees available for trap placement (Figures 5-6). Timed visual search became increasingly effective as the season progressed. In contrast, both BugBarrier and circle traps showed U-shaped seasonal patterns. Early in the season, tree-based traps outperformed timed visual search. By the end of the season, all methods had strong performance, though both tree-based traps again outperformed visual search on larger trees.

BugBarrier detection was higher on *A. altissima* than the other tree species surveyed. These differences were especially important for plots with very low *L. delicatula* density when the probability of detecting at least one individual was low and the choice of survey method could therefore determine if a population was detected. Thus, no single method consistently outperformed the other methods under all conditions.

The observed seasonal patterns in detection rates align with expectations based on *L. delicatula* biology. Timed visual search became more effective as the season progressed, consistent with the substantial increase in body size and conspicuousness that occurs as *L. delicatula* develops from small, black, early-instar nymphs to larger, red, late-instar nymphs and even larger adults.^57,58^ In contrast, detection by both tree-based traps declined during the middle of the season before increasing again later in the year. Both tree-based traps exploit the same tree climbing behavior pattern. The early season success of tree-based traps may be due in part to higher rates of falling and reclimbing for early instars^59^. The end-of season increase in traps’ effectiveness may be due in part to narrowing host preferences.^40,60^ Late-season improvements may also be driven by behavioral changes, including feeding directly on the trunks of some trees, mate-seeking behaviors, and oviposition. Together, our results indicate that detectability changed seasonally beyond what could be explained by changes in local abundance alone. Developmental stage, movement behavior, and host use preferences also likely influenced the effectiveness of survey methods.

The hierarchical Bayesian modelling framework employed here allowed us to estimate detection probabilities while accounting for two underlying ecological processes: whether a site was occupied during a given survey interval and, when occupied, how local abundance changed through time. The latent occupancy state of each site was allowed to change between sampling intervals. Conditional on occupancy, a constrained random walk allowed abundance within occupied sites to vary among successive sampling intervals as *L. delicatula* individuals dispersed into and from plots and suffered mortality within seasons (eqs. 1-7). This flexible structure accommodated substantial changes in *L. delicatula* densities over time, allowing for colonization and abundance fluctuations like those observed in the field.^38,61^ Recording counts rather than only presence/absence of *L. delicatula* within sites provided additional information about variation in local abundance, which can strongly influence the probability that a survey detects a population.^62,63^ The dynamic N-mixture approach therefore allowed us to estimate abundance-dependent detection probabilities rather than treating all occupied sites as equally detectable. This distinction is valuable for delimiting recent, low-density outbreaks and for tracking the effects of management treatments that substantially reduce *L. delicatula* numbers.^31,64^

Differences in the temporal scales of the sampling methods (5-minute timed search vs. 2-week trap deployment) required us to model abundance as a superpopulation rather than the straightforward number of individuals that are physically present at a site at the time of sampling.^65^ Specifically, the superpopulation represented all individuals that used a site at some point during a given approximately two-week sampling interval. Tree-based traps could encounter these individuals throughout the interval, whereas a five-minute visual search sampled only those members of the superpopulation that were present and available for observation during the brief search period. Consequently, estimated detection probabilities for timed visual search combine two processes: the probability that an individual in the two-week superpopulation was available within the plot during the search and the probability that an available individual was detected. This formulation was necessary to account for populations’ lack of closure over the two-week observation period. Future work with additional sampling and alternative sampling designs may be able to disentangle these two processes (availability and detection) to provide more information about visual search performance.

Our model also treated observations from the three survey methods as separate beta-binomial draws from the same underlying superpopulation rather than explicitly modeling removal of individuals by traps. A multinomial removal formulation^66^ could, in principle, account for individuals becoming unavailable for subsequent surveys after capture. However, such a formulation would require simultaneous observations from both trap types. Because circle traps were deployed later than BugBarrier traps during the first study year, this requirement would have resulted in discarding a substantial number of otherwise informative BugBarrier observations. The selected formulation therefore allowed us to retain more of the available data while integrating information across multiple survey methods.

As with other N-mixture model formulations, numerically separating abundance (N) from detection probability (p) presents an important inferential challenge.^67^ A relatively large population that is difficult to detect can generate observed counts similar to those from a small population that is easily detected. The beta-binomial observation models used here to account for overdispersion can further increase numerical trade-offs.^49^ We therefore assessed identifiability using data-cloning.^56^ Posterior variances decreased approximately in proportion to 1/*K* as the number of data clones (1,3,6) increased, indicating that these parameters were identifiable under our sampling design, if slow to mix. Specifically, our replication within the timed-visual surveys and the simultaneous use of multiple survey methods likely provided the needed additional information for separating abundance from detection.^67,68^

The three survey methods we compared are components of a broader toolset available for *L. delicatula* nymphs and adults. Terrestrial eDNA sampling ^69^ has demonstrated high sensitivity for detecting low-density populations and, in field comparisons, has detected *L. delicatula* more frequently than visual search.^33^ Previous studies on survey traps have likewise shown that tree species, trap design, and deployment conditions can strongly affect capture and detection.^37,38,70,71^ For example, in a prior comparison, circle traps captured more adult *L. delicatula* than BugBarrier traps, but the two traps did not differ in the probability of recording at least one detection. That study was deployed in heavily infested conditions where most traps captured at least one individual regardless of trap type, limiting the ability to distinguish the methods based on detection alone.^72^ During winter, surveillance shifts to target egg masses, which can be detected through visual surveys or trained detection dogs.^19,73–75^ Thus, these approaches complement one another by providing options across different life stages. Because previous studies differed in sampling design, population density, and the observation process being modeled, reported detection probabilities are not directly comparable. Future evaluations that deploy multiple survey methods simultaneously with standardized protocols and across shared gradients of abundance and season would provide more direct comparisons of their relative effectiveness.

Our results provide a basis for matching survey methods to field conditions rather than relying on a single approach throughout the growing season. When suitable large *A. altissima* are available, BugBarrier traps often provide the highest detection probabilities. During the midseason, and where suitable trees for trapping are unavailable, timed visual search can provide a competitive alternative. Circle traps may also perform well under particular combinations of tree size, tree species, season, and abundance, although they were less frequently the highest-performing method in our comparisons. Importantly, all three methods can have low detection probabilities when *L. delicatula* populations are small. Repeated surveillance and alternative methods like eDNA surveys may therefore be necessary. Detection probability is only one component of survey design, and managers must also consider labor requirements, material costs, tree availability, deployment time, and other logistical constraints. Nevertheless, incorporating realistic estimates of detection probability can improve both the allocation of survey effort and the interpretation of negative survey results. For *L. delicatula*, accounting for variation in detection across season, abundance, survey method, and trap-tree characteristics can help managers select survey approaches better matched to local conditions and improve the likelihood of detecting low-density populations when surveillance information is most consequential.

## Acknowledgements

We thank Nadia Garzione, David Long, Osariyekemwen Uyi, Elizabeth Wagner, and Brian Walsh for assistance in the field. We appreciate the help of Roana Fuller with modifying traps. We also thank all property owners for granting access to study locations. Funding was provided by Cooperative Agreements AP19PPQS&T00C080, AP21PPQS&T00C121 and AP24PPQS&T00C136 from the United States Department of Agriculture’s Animal and Plant Health Inspection Service (APHIS). The findings and conclusions in this publication are those of the author(s) and should not be construed to represent any official USDA or U.S. Government determination or policy.

