## Supplemental material for "Comparing pest survey methods through detection probability: seasonal and abundance-dependent detection of spotted lanternfly (*Lycorma delicatula*, Hemiptera: Fulgoridae)"

Keller, J. A., M. R. Helmus, K. Hoover. Detection of spotted lanternflies (*Lycorma delicatula*, Hemiptera: Fulgoridae) using tree-based traps and timed visual search varied seasonally and with abundance. Supplemental Material.

**Table S1**: Predictors included in the evaluated models for the two trap-based survey methods.

| Model | GDD | $GDD^2$ | DBH | TOH | Site RE |
|---|---|---|---|---|---|
| T1 | X | | X | X | |
| T2 | X | X | X | X | |
| T3 | X | | X | X | X |
| T4 | X | X | X | X | X |

**Table S2**: Predictors included in the evaluated models for the timed visual searches.

| Model | GDD | $GDD^2$ | log(N) | Observer RE |
|---|---|---|---|---|
| V1 | X | | | X |
| V2 | X | X | | X |
| V3 | X | | X | X |
| V4 | X | X | X | X |

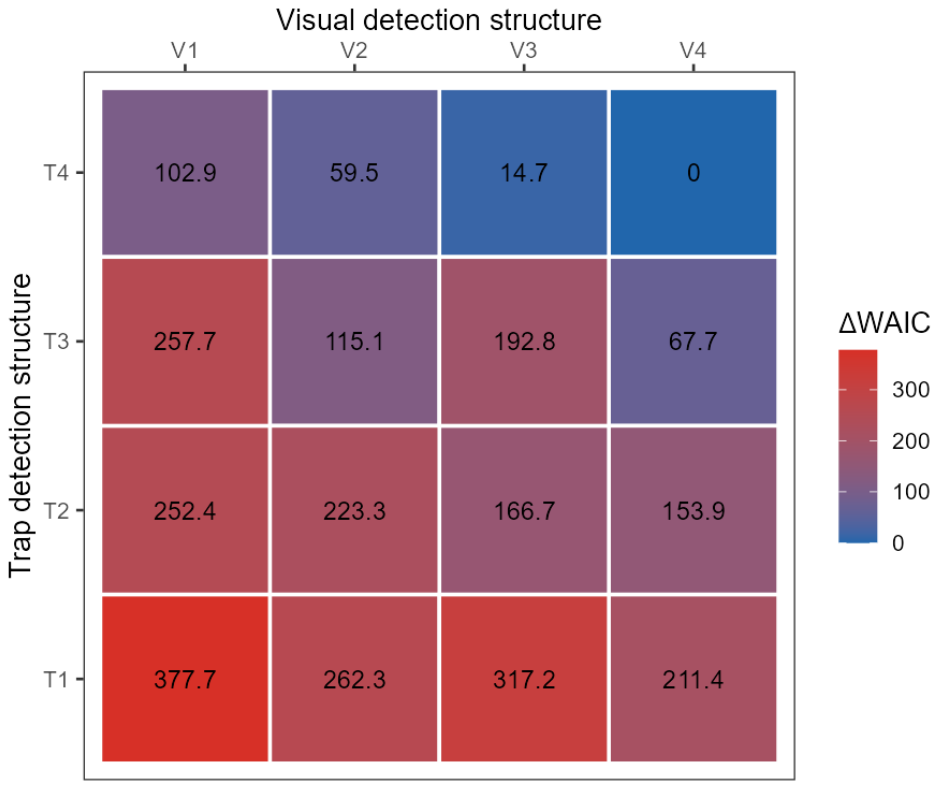


**Figure S1**: The full model including all assessed predictors for visual and trap-based survey methods had greater predictive performance than reduced models, as measured by WAIC. Tiles are colored and labeled according to the difference in WAIC relative to the lowest measured WAIC value (V4T4, see tables above for included predictors).

**Table S3:** Fitted model parameters

| Parameter | Description | Prior | Mean | Std. dev. | 2.50% | 50% | 97.50% | Rhat |
|---|---|---|---|---|---|---|---|---|
| $\psi$ | initial occupancy probability | Uniform(0, 1) | 0.649 | 0.056 | 0.536 | 0.650 | 0.758 | 1.0120 |
| $\gamma$ | colonization probability | Uniform(0, 1) | 0.070 | 0.019 | 0.038 | 0.069 | 0.111 | 1.0076 |
| $\varphi$ | persistence probability | Uniform(0, 1) | 0.998 | 0.002 | 0.992 | 0.998 | 1.000 | 1.0057 |
| ${\sigma_{abund}}^2$ | abundance random walk standard deviation | Uniform(0, 3) | 0.653 | 0.030 | 0.595 | 0.653 | 0.715 | 1.0008 |
| $\beta_{visual_0}$ | visual intercept | Normal(mean = 0, SD = 10) | -3.116 | 0.255 | -3.630 | -3.109 | -2.632 | 1.0066 |
| $\beta_{visual_{GDD}}$ | visual linear GDD effect | Normal(mean = 0, SD = 10) | 0.582 | 0.065 | 0.457 | 0.581 | 0.709 | 1.0179 |
| $\beta_{visual_{GDD^2}}$ | visual squared GDD effect | Normal(mean = 0, SD = 10) | -0.178 | 0.064 | -0.305 | -0.179 | -0.050 | 1.0073 |
| $\beta_{visual_{\log(N)}}$ | visual N dependent detection | Normal(mean = 0, SD = 10) | 0.403 | 0.047 | 0.314 | 0.402 | 0.497 | 1.0064 |
| $\varepsilon_{observer_1}$ | visual observer effect 1 | Normal(mean = 0, SD = 2) | -0.365 | 0.069 | -0.499 | -0.366 | -0.228 | 1.0006 |
| $\varepsilon_{observer_2}$ | visual observer effect 2 | Normal(mean = 0, SD = 2) | 0.036 | 0.064 | -0.087 | 0.035 | 0.165 | 1.0003 |
| $\varepsilon_{observer_3}$ | visual observer effect 3 | Normal(mean = 0, SD = 2) | 0.289 | 0.059 | 0.177 | 0.287 | 0.405 | 1.0005 |
| $\varepsilon_{observer_4}$ | visual observer effect 4 | Normal(mean = 0, SD = 2) | 0.206 | 0.062 | 0.086 | 0.205 | 0.331 | 1.0005 |
| $\varepsilon_{observer_5}$ | visual observer effect 5 | Normal(mean = 0, SD = 2) | 0.173 | 0.212 | -0.251 | 0.178 | 0.577 | 1.0005 |
| $\varepsilon_{observer_6}$ | visual observer effect 6 | Normal(mean = 0, SD = 2) | -0.339 | 0.167 | -0.679 | -0.334 | -0.017 | 1.0001 |
| $\rho_{visual}$ | beta-binomial concentration for visual | Gamma(shape = 5, rate = 0.2); average value ≈ 25 | 35.418 | 5.722 | 25.054 | 35.137 | 47.206 | 1.0022 |
| $\beta_{trap_{0_1}}$ | Bugbarrier intercept | Normal(mean = 0, SD = 10) | -3.937 | 0.205 | -4.343 | -3.938 | -3.534 | 1.0007 |
| $\beta_{trap_{GDD_1}}$ | Bugbarrier linear GDD effect | Normal(mean = 0, SD = 10) | 0.180 | 0.060 | 0.063 | 0.180 | 0.296 | 1.0132 |
| $\beta_{trap_{GDD^2_1}}$ | Bugbarrier squared GDD effect | Normal(mean = 0, SD = 10) | 0.666 | 0.064 | 0.542 | 0.666 | 0.792 | 1.0055 |
| $\beta_{trap_{DBH_1}}$ | Bugbarrier DBH effect | Normal(mean = 0, SD = 10) | 0.307 | 0.057 | 0.196 | 0.306 | 0.419 | 1.0005 |
| $\beta_{trap_{TOH_1}}$ | Bugbarrier tree of heaven effect | Normal(mean = 0, SD = 10) | 0.496 | 0.175 | 0.154 | 0.496 | 0.840 | 1.0002 |
| ${\sigma_{site_1}}^2$ | Bugbarrier site random effect standard deviation | Precision ~ Gamma(shape = 0.01, rate = 0.01) | 0.598 | 0.083 | 0.449 | 0.593 | 0.774 | 1.0003 |
| $\rho_{trap_1}$ | beta-binomial concentration for Bugbarrier | Uniform(0.1, 100) | 14.803 | 2.482 | 10.458 | 14.626 | 20.117 | 1.0014 |
| $\beta_{trap_{0_2}}$ | circle trap intercept | Normal(mean = 0, SD = 10) | -3.835 | 0.230 | -4.293 | -3.835 | -3.389 | 1.0007 |
| $\beta_{trap_{GDD_2}}$ | circle trap linear GDD effect | Normal(mean = 0, SD = 10) | 0.142 | 0.064 | 0.018 | 0.142 | 0.266 | 1.0100 |
| $\beta_{trap_{GDD^2_2}}$ | circle trap squared GDD effect | Normal(mean = 0, SD = 10) | 0.343 | 0.071 | 0.203 | 0.343 | 0.480 | 1.0088 |
| $\beta_{trap_{DBH_2}}$ | circle trap DBH effect | Normal(mean = 0, SD = 10) | 0.313 | 0.068 | 0.180 | 0.313 | 0.447 | 1.0005 |
| $\beta_{trap_{TOH_2}}$ | circle trap tree of heaven effect | Normal(mean = 0, SD = 10) | 0.190 | 0.209 | -0.224 | 0.191 | 0.600 | 1.0001 |
| ${\sigma_{site_2}}^2$ | circle trap site random effect standard deviation | Precision ~ Gamma(shape = 0.01, rate = 0.01) | 0.683 | 0.101 | 0.503 | 0.677 | 0.897 | 1.0004 |
| $\rho_{trap_2}$ | beta-binomial concentration for circle trap | Uniform(0.1, 100) | 15.237 | 2.671 | 10.602 | 15.040 | 21.016 | 1.0008 |

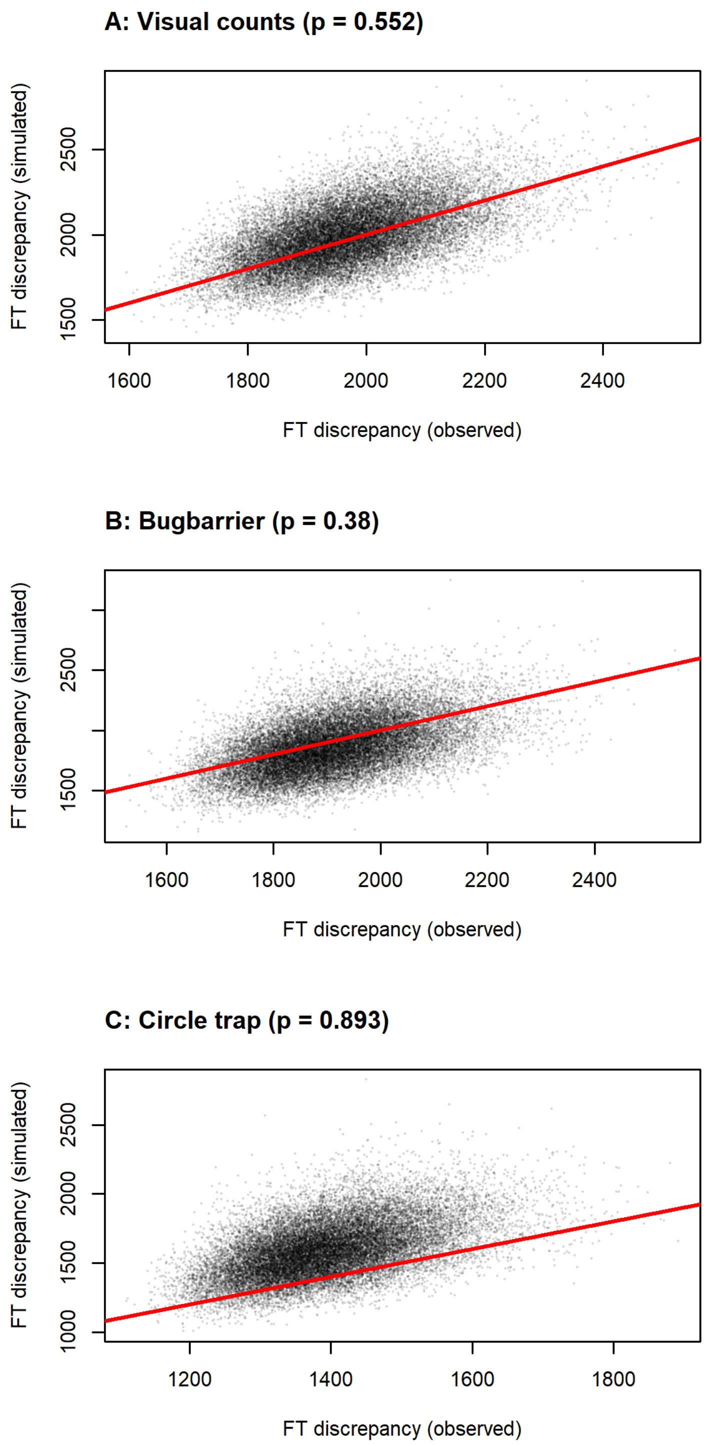


**Figure S2**: Freeman-Tukey discrepancy results for (A) timed visual search, (B) Bugbarrier, and (C) circle trap observations. The 1:1 line is shown in red within each panel. Bayesian p-values for these model components were 0.55, 0.38, and 0.89 respectively.

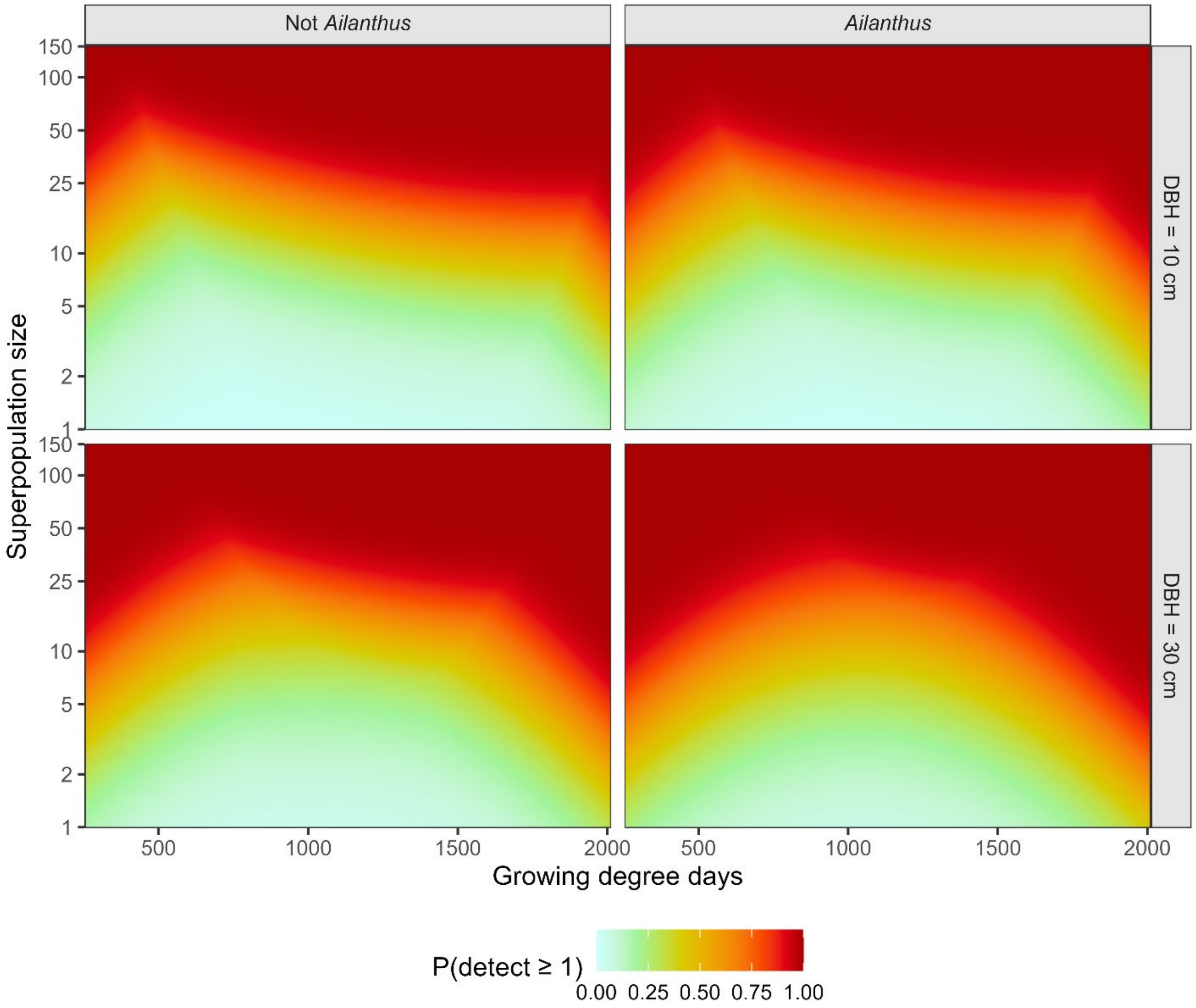


**Figure S3**: The highest mean probability of detection that can be achieved by selecting the best of the evaluated methods under various conditions. Panels show estimated maximum mean detection probability for tree of different sizes (rows), species (columns) and at differing superpopulation sizes (y-axes in each panel) and seasonal timings (cumulative growing degree days along the x-axis in each panel).

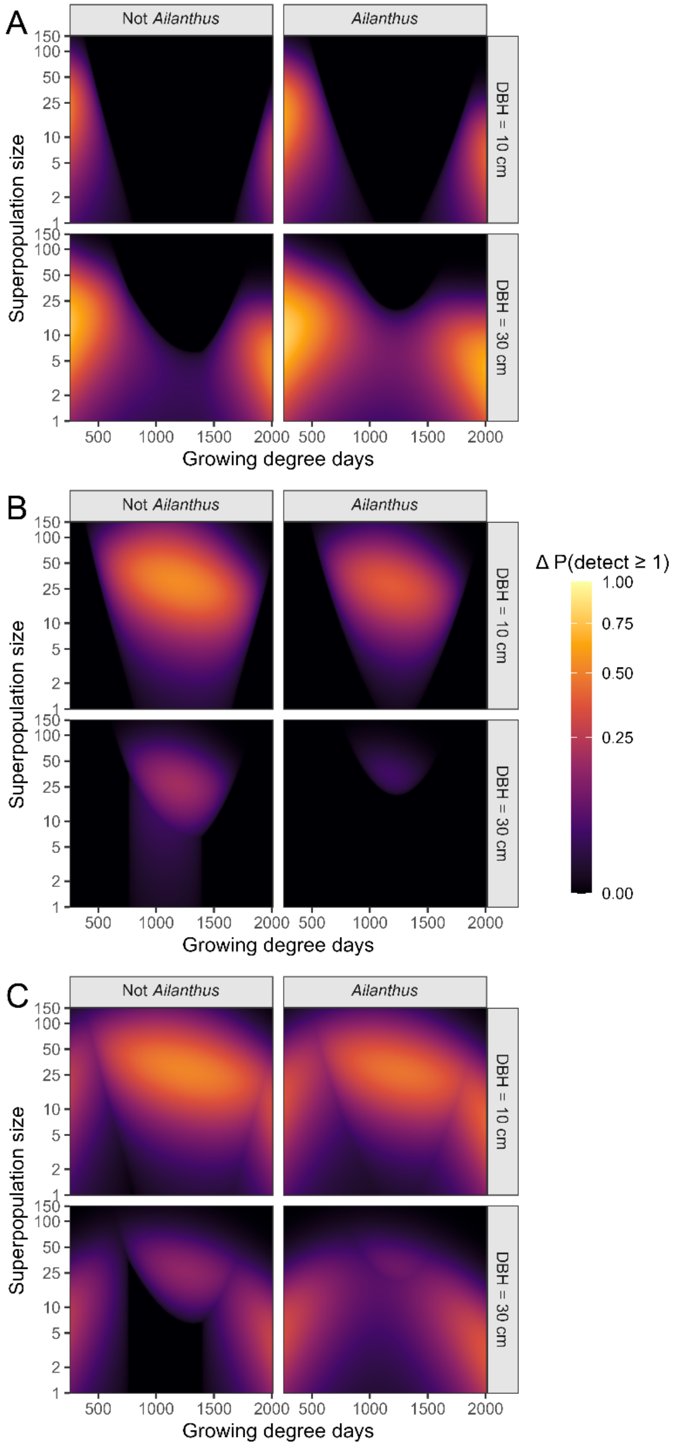


**Figure S4:** Difference between the detection probability of the best-performing method and that of (A) timed visual search (B) Bugbarrier traps and (C) circle traps. Methods are compared on different host plants (panel columns) tree sizes (panel rows), seasonal timings (x axis within panels) and *L. delicatula* densities (y axis within panels).

**Table S4**: Data-cloning results.[†]

| K | Parameter | Mean | Standard deviation ($SD_K$) | Standard deviation with K = 1 ($SD_1$) | $R_K$ |
|---|---|---|---|---|---|
| 3 | beta.vis.0 | -3.14 | 0.15 | 0.25 | 1.15 |
| 3 | beta.vis.gdd | 0.59 | 0.04 | 0.06 | 1.13 |
| 3 | beta.vis.gdd2 | -0.18 | 0.04 | 0.07 | 0.97 |
| 3 | beta.vis.logN | 0.40 | 0.03 | 0.04 | 1.19 |
| 3 | rho.vis | 37.07 | 3.80 | 5.62 | 1.37 |
| 3 | gamma.obs[1] | -0.37 | 0.04 | 0.07 | 0.96 |
| 3 | gamma.obs[2] | 0.03 | 0.04 | 0.06 | 0.93 |
| 3 | gamma.obs[3] | 0.28 | 0.03 | 0.06 | 0.99 |
| 3 | gamma.obs[4] | 0.20 | 0.04 | 0.06 | 0.97 |
| 3 | gamma.obs[5] | 0.18 | 0.12 | 0.21 | 0.96 |
| 3 | gamma.obs[6] | -0.33 | 0.09 | 0.17 | 0.95 |
| 3 | beta.bb.0 | -3.95 | 0.12 | 0.20 | 1.00 |
| 3 | beta.bb.gdd | 0.18 | 0.04 | 0.06 | 1.05 |
| 3 | beta.bb.gdd2 | 0.66 | 0.04 | 0.06 | 1.01 |
| 3 | beta.bb.dbh | 0.31 | 0.03 | 0.06 | 1.01 |
| 3 | beta.bb.toh | 0.50 | 0.10 | 0.17 | 0.96 |
| 3 | sigma.bb.site | 0.58 | 0.05 | 0.08 | 0.96 |
| 3 | rho.bb | 14.78 | 1.40 | 2.37 | 1.05 |
| 3 | beta.ct.0 | -3.85 | 0.13 | 0.23 | 1.01 |
| 3 | beta.ct.gdd | 0.15 | 0.04 | 0.06 | 1.03 |
| 3 | beta.ct.gdd2 | 0.34 | 0.04 | 0.07 | 1.01 |
| 3 | beta.ct.dbh | 0.31 | 0.04 | 0.07 | 1.00 |
| 3 | beta.ct.toh | 0.20 | 0.12 | 0.21 | 0.98 |
| 3 | sigma.ct.site | 0.66 | 0.06 | 0.10 | 0.91 |
| 3 | rho.ct | 15.16 | 1.51 | 2.59 | 1.02 |
| 3 | psi | 0.66 | 0.04 | 0.06 | 1.12 |
| 3 | gamma | 0.07 | 0.01 | 0.02 | 0.98 |
| 3 | phi | 1.00 | 0.00 | 0.00 | 0.33 |
| 3 | sigma.proc | 0.66 | 0.02 | 0.03 | 1.11 |
| 6 | beta.vis.0 | -3.20 | 0.10 | 0.25 | 0.93 |
| 6 | beta.vis.gdd | 0.59 | 0.03 | 0.06 | 1.07 |
| 6 | beta.vis.gdd2 | -0.18 | 0.03 | 0.07 | 1.03 |
| 6 | beta.vis.logN | 0.40 | 0.02 | 0.04 | 0.94 |
| 6 | rho.vis | 38.96 | 2.37 | 5.62 | 1.07 |
| 6 | gamma.obs[1] | -0.37 | 0.03 | 0.07 | 0.95 |
| 6 | gamma.obs[2] | 0.03 | 0.03 | 0.06 | 0.92 |
| 6 | gamma.obs[3] | 0.28 | 0.02 | 0.06 | 0.96 |

| K | Parameter | Mean | Standard deviation ($SD_K$) | Standard deviation with K = 1 ($SD_1$) | $R_K$ |
|---|---|---|---|---|---|
| 6 | gamma.obs[4] | 0.20 | 0.02 | 0.06 | 0.94 |
| 6 | gamma.obs[5] | 0.18 | 0.08 | 0.21 | 0.93 |
| 6 | gamma.obs[6] | -0.33 | 0.07 | 0.17 | 0.92 |
| 6 | beta.bb.0 | -3.98 | 0.08 | 0.20 | 0.90 |
| 6 | beta.bb.gdd | 0.19 | 0.02 | 0.06 | 0.97 |
| 6 | beta.bb.gdd2 | 0.66 | 0.03 | 0.06 | 0.97 |
| 6 | beta.bb.dbh | 0.31 | 0.02 | 0.06 | 0.97 |
| 6 | beta.bb.toh | 0.49 | 0.07 | 0.17 | 0.96 |
| 6 | sigma.bb.site | 0.58 | 0.03 | 0.08 | 0.91 |
| 6 | rho.bb | 15.35 | 0.96 | 2.37 | 0.98 |
| 6 | beta.ct.0 | -3.89 | 0.09 | 0.23 | 0.90 |
| 6 | beta.ct.gdd | 0.15 | 0.03 | 0.06 | 0.99 |
| 6 | beta.ct.gdd2 | 0.34 | 0.03 | 0.07 | 1.00 |
| 6 | beta.ct.dbh | 0.31 | 0.03 | 0.07 | 0.97 |
| 6 | beta.ct.toh | 0.20 | 0.08 | 0.21 | 0.95 |
| 6 | sigma.ct.site | 0.65 | 0.04 | 0.10 | 0.90 |
| 6 | rho.ct | 15.69 | 1.05 | 2.59 | 0.98 |
| 6 | psi | 0.66 | 0.02 | 0.06 | 1.05 |
| 6 | gamma | 0.06 | 0.01 | 0.02 | 0.94 |
| 6 | phi | 1.00 | 0.00 | 0.00 | 0.20 |
| 6 | sigma.proc | 0.66 | 0.01 | 0.03 | 0.96 |

† K indicates the number of identical clones of the dataset combined in the likelihood. $R_K = SD_K \times \sqrt[2]{K} / SD_1$ is the standardized posterior standard deviation, which converges to 1 as K increases if a parameter is identifiable.
